# A structured scaffold underlies activity in the hippocampus

**DOI:** 10.1101/2021.11.20.469406

**Authors:** Dounia Mulders, Man Yi Yim, Jae Sung Lee, Albert K. Lee, Thibaud Taillefumier, Ila R. Fiete

## Abstract

Place cells are believed to organize memory across space and time, inspiring the idea of the cognitive map. Yet unlike the structured activity in the associated grid and head-direction cells, they remain an enigma: their responses have been difficult to predict and are complex enough to be statistically well-described by a random process. Here we report one step toward the ultimate goal of understanding place cells well enough to predict their fields. Within a theoretical framework in which place fields are derived as a conjunction of external cues with internal grid cell inputs, we predict that even apparently random place cell responses should reflect the structure of their grid inputs and that this structure can be unmasked if probed in sufficiently large neural populations and large environments. To test the theory, we design experiments in long, locally featureless spaces to demonstrate that structured scaffolds undergird place cell responses. Our findings, together with other theoretical and experimental results, suggest that place cells build memories of external inputs by attaching them to a largely prespecified grid scaffold.

## Introduction

The hippocampus plays an essential role in episodic memory [1, 2, 3] and more generally in organizing memory across space and time [4, 5, 6]. It is believed to construct cognitive maps that link the “what” and the “when” to the “where” of experience to enable rich contextual inference based on memory [5, 7]. In line with this view, place cells encode a combination of variables including spatial location, sensory cues [8, 9], and abstract non-spatial variables including time [10, 11].

However, unlike their spatially structured counterparts such as head direction and grid cells – for which we possess good models [12, 13, 14] with detailed experimental validations of these models [15, 16, 17, 18, 19, 20] and thus a good explanation of the circuit mechanisms underlying their function as velocity integrators and analog memories – it is unclear how the hippocampus and place cells encode the cognitive maps that link *what* to *where* in memory.

When recorded in small spatial environments, place cells have simple response properties, with each cell exhibiting a localized field in the neighborhood of some location in the explored 2D space [21]. However, the full reality is more complex: hippocampal cells transition from place coding to ensemble coding in large environments, with each cell exhibiting multiple fields [22, 23]. The coding scheme that governs field placement has been difficult to decipher, and the best quantification of place field statistics so far is as a random process [23].

Recent theoretical work indicates that mnemonic place fields (i.e., fields that may be relatively reliably activated even without a localized external cue at the position of the field) of individual cells cannot be placed at arbitrary locations and must be highly constrained in their organization [24]. Innovative empirical work to induce a new mnemonic place field at some arbitrarily chosen location through spatially targeted current injection into the cell reveals that while it is possible to do so, the precise position is not under experimental control [25]– the cell ultimately selects a location based presumably on some internal constraints [26], and moreover this induction results in the appearance and disappearance of additional, untargeted fields [27], also consistent with recent predictions [24]. These findings suggest that (mnemonic or non-cued) place field configurations are bound by some specific constraints; our goal in this work is to unmask the nature of this structure.

Here, within a specific theoretical framework for the origins of the CA1 place cell response, we predict a strong spatial scaffold in place cells, inherited from grid cells. We then test this prediction by performing large-scale neural recordings in large locally featureless spaces to show for the first time the existence of a regular organization in place fields. The finding is consistent with our theoretical analyses in which we show that though a scaffold should not be visible in single cells it will be unmasked in the population response. We conclude that the collective place cell population generates, with the help of multi-modular grid cell inputs, a regular and predetermined coding structure for the organization of memory [28, 29, 3, 27]. This organization of states might permit a much larger memory capacity than suggested by standard autoassociative models of hippocampus that do not incorporate such structure [24].

## Results

### A model of place cells undergirded by feedforward grid cells reproduces the random statistics of place cells in large environments

We espouse a theoretical framework in which place fields are driven by a combination of grid inputs and external spatial cues derived from landmarks, borders, and other features in the environment, Fig. 1a. In this framework, grid cells generate velocity-derived positional estimates and are are a primary driver of mnemonic place fields, which are defined as fields not primarily driven by localized external cues, Fig. 1b,d. The model also includes noisy inputs that are spatially unreliable. This framework contrasts with the one in which grid cells are primarily a downstream consequence of place cell activity, Fig. S1b [30, 31, 32]. All these inputs project sparsely through random log-normally distributed weights [33] to place cells, which apply a threshold to generate fields (Methods). The threshold may be viewed as determined individually by neurons or influenced by local recurrent inhibition.

**Fig. 1:**
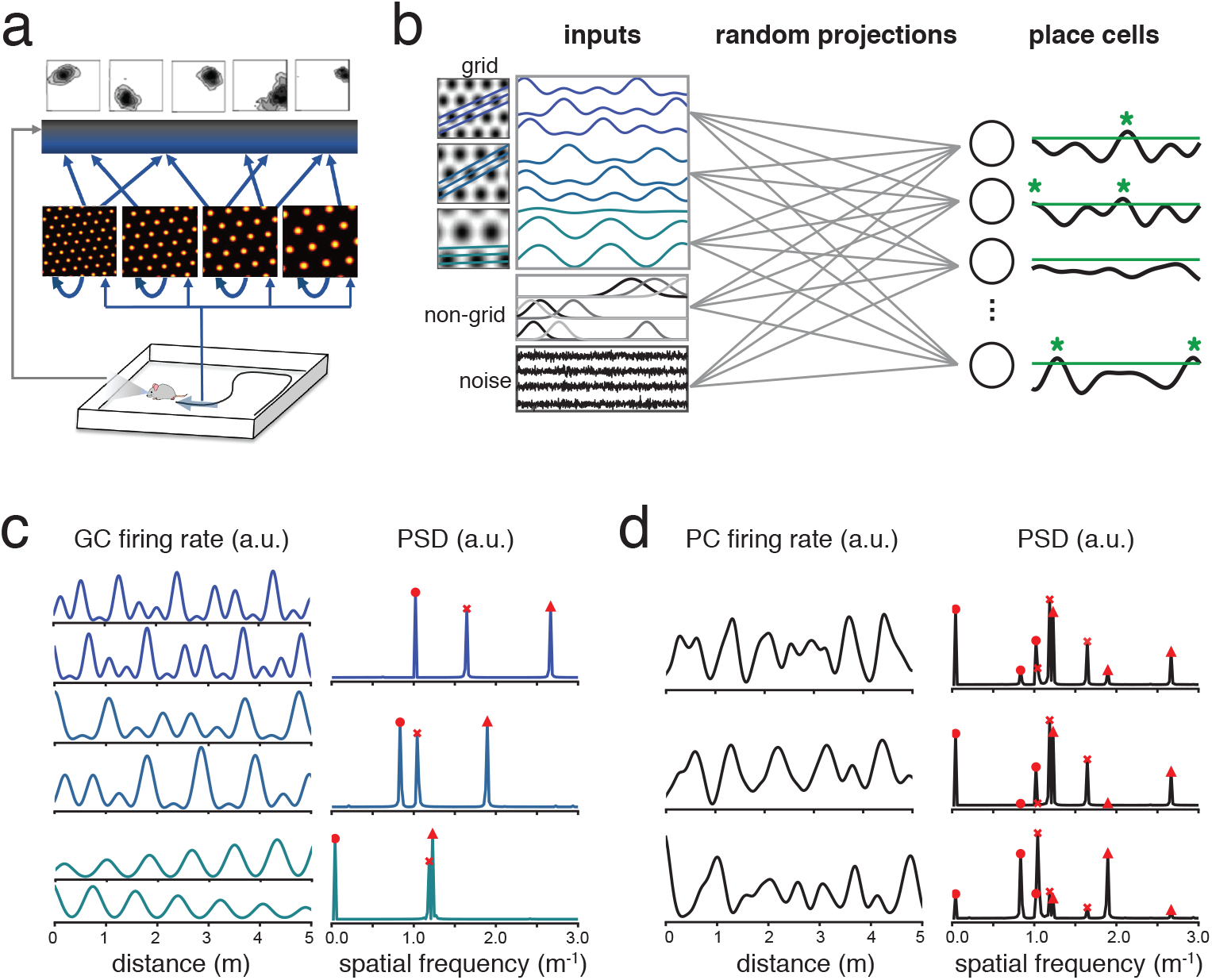
Grid-driven place cell model theoretically predicts structure in Fourier space. **a**, Theoretical framework positing that grid cells provide a primary internally generated motion integration-based portion of the spatial tuning of CA1 place cells, which is combined with externally derived cues to form place fields. **b**, A population of place cells (PCs) in a 1D environment receives inputs from grid cells (GCs; 1D grid responses are modeled as slices through spatially periodic 2D lattices [36]) from different modules; spatially tuned non-periodic inputs (such as from external cues) from non-grid neurons; and spatially untuned inputs modeled as noise. A threshold (green line) is applied to PC inputs to define place fields (green asterisks). **c**, Responses of GCs in 1D are characterized by three main power spectral density (PSD) peaks (indicated with red markers), whose positions depend on the underlying 2D grid period and orientation of the 1D slice [36]. Different colors indicate distinct grid modules; co-modular cells express the same set of PSD peaks. **d**, Prediction of structure in place cell PSDs: If PC activities were obtained through random projections of grid inputs only, their PSDs would consist of a weighted sum of the peaks from all of the PSDs of the constituent GC modules in c. The relative amplitudes of these peaks across PCs depend on the random mappings. For conceptual clarity, the number of GCs projecting to each PC here is small and thresholds are low, which makes PC fields non-sparse but PSDs sparse; realistic values are used in Fig. 2, which generate sparse fields and dense PSDs.)

Despite its simplicity, the model exhibits a number of interesting features. First, the responses of model place cell populations resemble recorded responses [23], appearing random and quantitatively reproducing the field statistics of place cells even when the only inputs are structured grid responses: the distribution of number of fields across cells is described by a negative-binomial (gamma-Poisson) distribution and the recruitment of fields along the track is memoryless, Fig. 2a. The inter-field interval distribution of all cells (normalized by the cell’s place field probability) follows an exponential distribution, as expected for a Poisson process. In sum, random-looking field statistics do not imply an absence of structure. Second, a single value of skewness in the log-normal input weight distribution is sufficient to capture multiple statistical measures of place cell activity, including the number of place fields over a given track length, the fraction of cells with a field, and the parameters of the gamma-Poisson (negative binomial) function that best fits the long-tailed distribution of fields per cell (SI Fig. S2d). Third, the weight distribution corresponding to the value of model skewness that reproduces experimental place cell statistics resembles measured weight distributions in the brain [33] (SI Fig. S2e). Fourth, model place cells exhibit persistent firing propensities across space: a cell that is highly (un)likely to fire on the first half of a long track is also highly (un)likely to do so later. The persistence of firing propensity differences in this model emerges from network-level properties, without the assumption of persistent biophysical differences in cellular excitability [34, 35]: the diversity of propensities is due to the heterogeneity of the weights between grid and place cells, while the persistence is because the weights are fixed and the activity level of each grid cell is sustained across space. Thus, a place cell model with structured grid inputs reproduces a number of known properties of place cells recorded on long tracks.

**Fig. 2:**
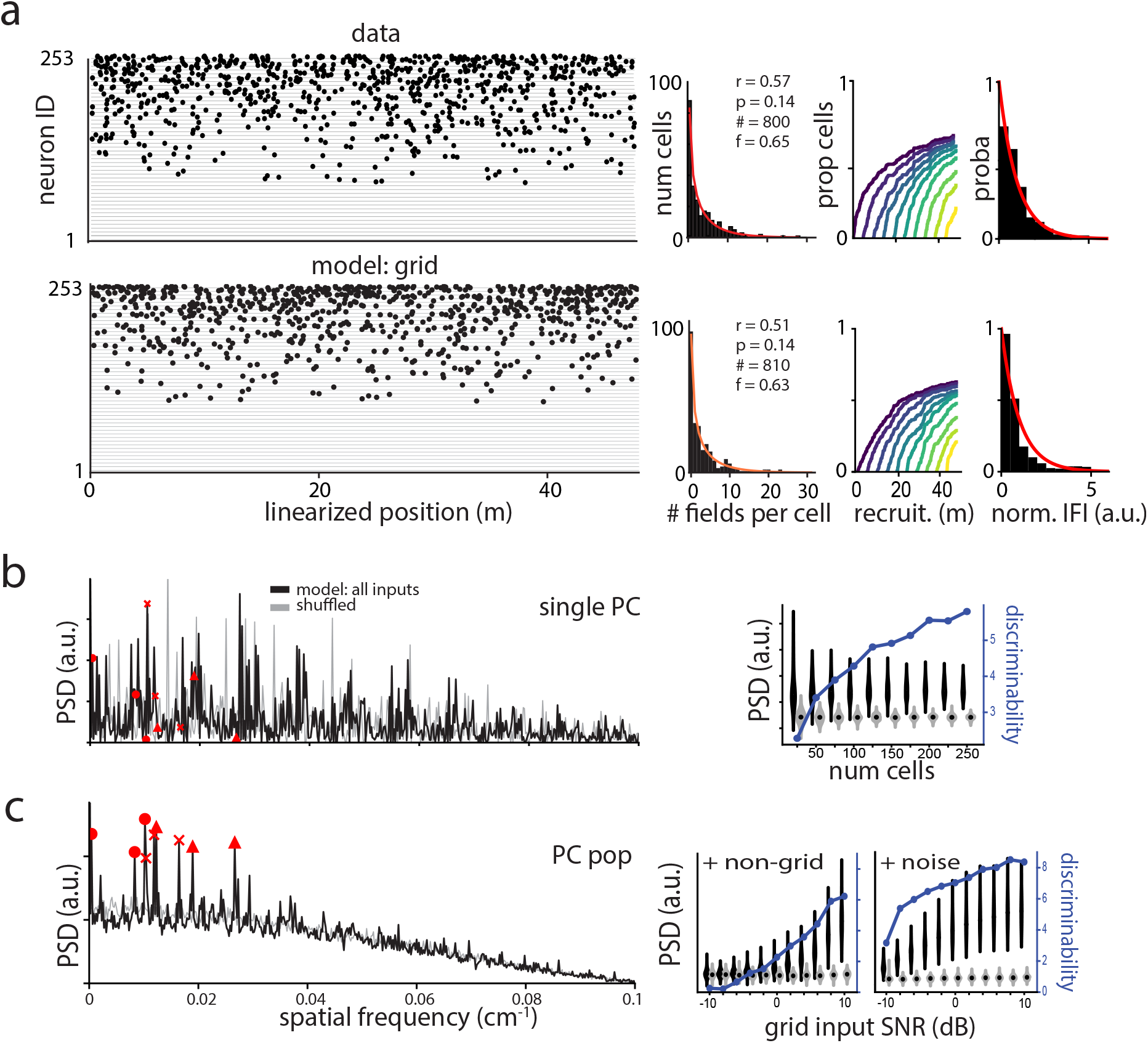
Model generates random-looking field locations consistent with data, with predicted structure detectable only after large-scale spectral averaging. **a**, Top: Place fields from electrophysiological recordings of 253 place cells [23], with quantification of population-level statistics (red curve: Gamma-Poisson distribution fit to data. r, p: Gamma-Poisson shape parameters, #: total number of place fields, f: fraction of cells having at least one field along the track (%)). The recruitment of place cells along the track is memoryless and inter-field intervals (IFIs) pooled across cells (for cells with at least 6 fields, after normalizing by mean rate of place fields per cell) are exponentially distributed (far right). Bottom: place fields of 253 place cells generated by our model (Methods) even when using only periodic grid cell inputs exhibit similar population-level Gamma-Poisson statistics and memoryless recruitment, and normalized IFIs pooled across cells (for cells with at least 6 fields) follow an exponential distribution. **b**, PSD of the place fields of a single model PC with grid cell (GC), non GC and noise inputs (GC to non GC and noise SNR = 10 dB). Red markers indicate positions of predicted peaks at the frequencies induced by the 3 GC input modules, with one marker for each spatial frequency (Methods). Right: Violin plots give the distributions of PSD amplitudes at these GC-related frequencies in averaged PSD as a function of the number of cells in the averages, using *n* = 30 simulations. The PSDs are compared to PSDs of shuffled responses. Blue curve: discriminability index between the shuffled and real distributions of amplitudes, computed using the average standard deviation [52] (Methods). **c**, Averaged PSD computed from the PSDs of 253 place cells with grid, non-grid and noise inputs (GC to non GC and noise SNR = 10 dB, leading to an average ratio of input GC to non GC activity bumps of 14.42, although each GC produces an average of 62.33 input bumps and each non GC a single bump, see Methods). Right: Violin plots indicate distributions of PSD amplitudes at GC-related frequencies with *n* = 30 simulations and as a function of the relative amplitude of non-grid to grid inputs, and the relative amount of noise to grid inputs.

### Predicted structure in large-scale spectral averages of place cells

Since the spatial responses of co-modular grid cells form a triangular lattice with common period and orientation, in Fourier space they are dominated by a common set of three primary components [36]. Though grid cells have non-periodic responses on linear 1D tracks, they are well-described by a slice through the same 2D lattice because they exhibit the same three primary Fourier components [36] (Fig. 1b-c, left). This result suggests that under perfectly idealized conditions – with place cells driven by only periodic inputs from a small number of grid cells, with no noise, external sensory cues, or path integration error – the power spectral density (PSD) of each place cell should exhibit all and only the spectral peaks of its grid inputs (Fig. 1c-d, right).

Under realistic conditions, is it possible to find an underlying structure in place cells? On long tracks, path integration errors degrade the periodicity of grid input and hence decrease the spectral SNR of the inputs (SI Fig. S3b). On the other hand, when tracks are limited in length, the ability to identify key PSD peaks of even ideal grid cells becomes limited [36]. Noise and non-grid inputs to place cells would further erode any grid-driven peaks in the place cell PSDs. For all these reasons, we do not theoretically expect individual place cells to reveal structure in their field arrangements (Fig. 2b and analytical derivation in SI) even if strongly driven by structured inputs, consistent with experimental findings to date.

However, grid-driven spectral peaks in the place cell responses have a unique feature relative to peaks from other types of input: they are shared across place cells that receive inputs from common sets of grid modules. This means that grid-driven peaks will sum coherently if we add PSDs across place cells, while noise-driven inputs and non-grid inputs, which are not as broadly shared, will not (Fig. S2b-c, and SI for analytical derivation). Thus, summing per-cell and per-lap

PSDs is predicted to produce distinguishable structure peaks (Fig. 2c, left and SI) that become increasingly discriminable for larger population sizes (Fig. 2b, right and Fig. S2b, right), even in the presence of large non-grid inputs (Fig. 2c, right). Path integration (PI) errors diminish the SNR of key structure peaks, but are predicted, even at very large values (corresponding to 1 cm s.d. per 1 cm displacement), to allow many of the key peaks to remain discriminable (SI Fig. S3b compared to Fig. S2b). This is because PSD peaks depend on place field separations between adjacent and otherwise nearby fields rather than on absolute field positions, and are thus not vulnerable to slowly accumulating PI errors within and across laps.

In sum, the primary prediction of the model is that spatial frequencies associated with grid cell firing should be robustly over-represented in the sum of the per-cell and per-lap power spectral densities of place cells, and this structure should become visible in large-scale population recordings on long tracks.

### Statistics of mnemonic place fields in large virtual environments match known field statistics

We designed a 40-meter linear virtual track (Fig. 3a-b) with irregularly and closely spaced sloping stripes on the walls that enable optic flow-based motion estimation but convey no spatial information, Fig. 3b. To minimize cue-driven responses and focus on mnemonic place fields, the only spatial features in the environment were large orienting cues at the start of the track or rendered far from the track. Two-photon calcium imaging in head-fixed mice, Fig. 3a, allowed us to simultaneously record ∼500 −1000 CA1 pyramidal cells. Animals were trained to run from one end of the track to the other with randomly dispensed liquid rewards, Fig. 3e. Place fields are defined from the fluorescence traces of cellular ROIs (Fig. 3c) by filtering followed by application of an amplitude and a duration threshold (Fig. 3d; see Methods and SI for details; note that place fields were defined per lap in this manner, because the absence - by design - of localized landmarks that usually correct spatial estimation drift means that absolute place field positions are generally not preserved across laps [37]).

**Fig. 3:**
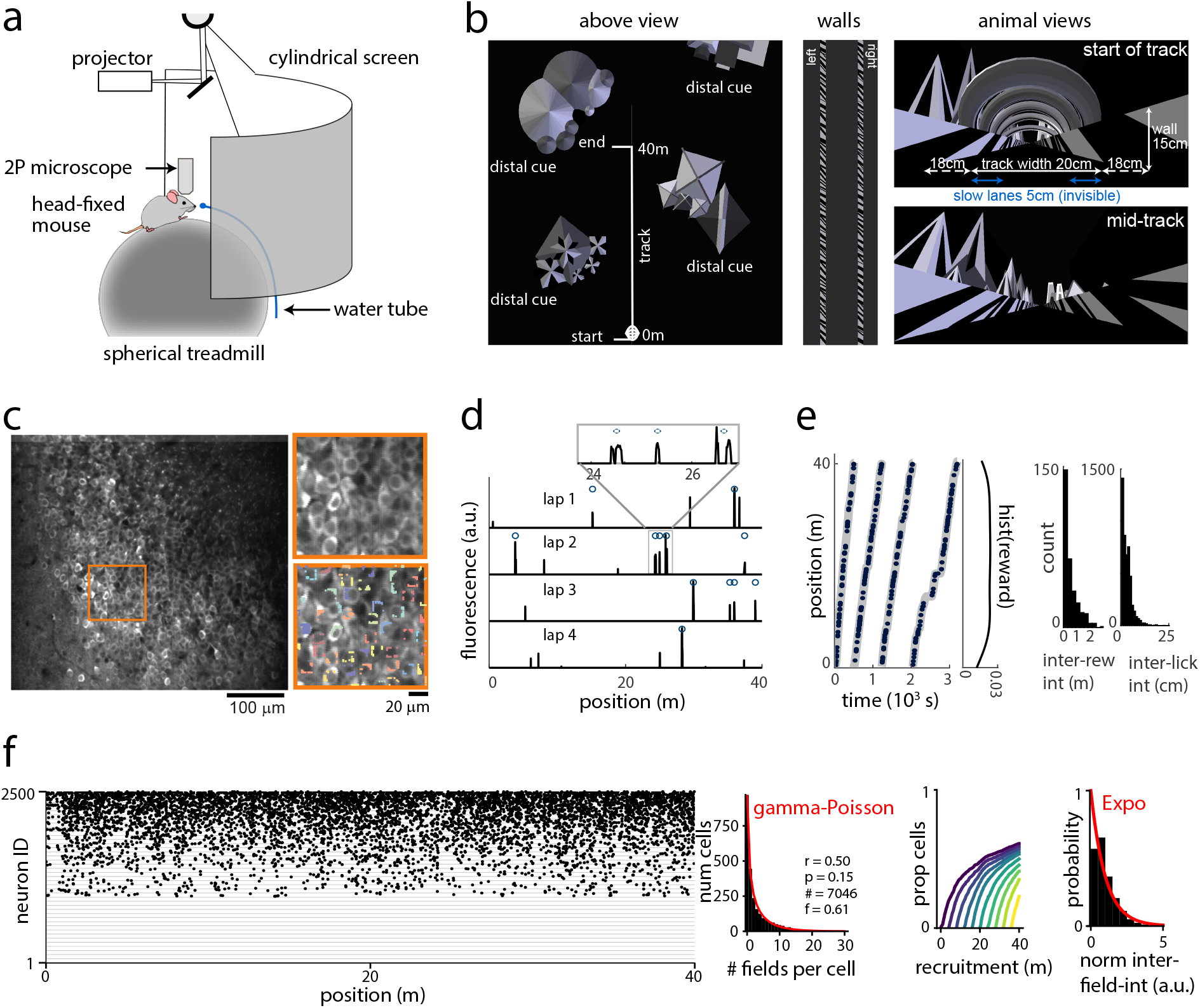
Place cell population responses in long landmark-free linear tracks. **a**, Experimental setup. **b**, Description of the linear track. There is a tunnel-like structure at the start of the track. The proximal walls patterns consist in black stripes with a tilt of 25, 35 or 45°, with inter-stripe intervals uniformly distributed between 2 and 20 cm (middle panel). There are two distal cues on each side of the track (big mountains). **c**, Fluorescence image of CA1 pyramidal neurons genetically expressing GCaMP6f and ROI of a small subset of cells (mouse AN11). **d**, Example of calcium traces from one place cell in mouse AN08. Dots indicate the detected place fields. Place fields are defined per lap because of the expected drift in spatial localization along long, locally featureless tracks (see Results and Methods). **e**, Trajectories of one mouse (AN08) across the 4 laps on the linear track (left). Dots indicate the reward positions, and the empirical reward probability is given in the right inset. Empirical distributions of the inter-reward and inter-licking intervals in AN08 (right). **f**, Distribution of the number of place fields in all the recorded CA1 cells (n = 2500, 4 mice). Fit of Gamma-Poisson distribution (red). r, p: Gamma-Poisson shape parameters, #: total number of fields, f: fraction of place cells having at least one place field. The recruitment of place cells is memoryless and the inter-field-intervals normalized per cell follow an exponential distribution, as expected for random field locations only characterized by a mean rate.

The basic statistics of place fields and field intervals in this large featureless environment, Figs. 3f and S4, match previous findings from cue-rich linear and non-linear tracks in real-space [23, 34] (cf. Fig. 2a). This suggests that though place cells can be influenced by reward location [38, 34] and sensory landmarks [9], the underlying statistical properties of place cell firing are largely independent of these features and may be driven by intrinsic mechanisms.

### Evidence of grid-driven scaffold in place cell population responses

We computed the population-level PSD by summing the per-cell and per-lap PSDs, which we obtained from the filtered raster of place field locations (see Methods). We identified significant peaks in the population PSD by applying a set of simple but stringent criteria. Peaks were first identified by the topological measure of prominence, Fig. 4a. The subset of prominent peaks that were also statistically significant relative to PSD peaks from shuffled data were identified, Fig. 4b. Next, we applied a consistency criterion: cells were repeatedly (and randomly) partitioned into four sets, Fig. 4c. A peak was deemed consistent if in 95% of the random partition repeats, the peak was present in ≥50% of the sets, Fig. 4d. The final set of selected peaks was thus prominent, significant, and consistent.

**Fig. 4:**
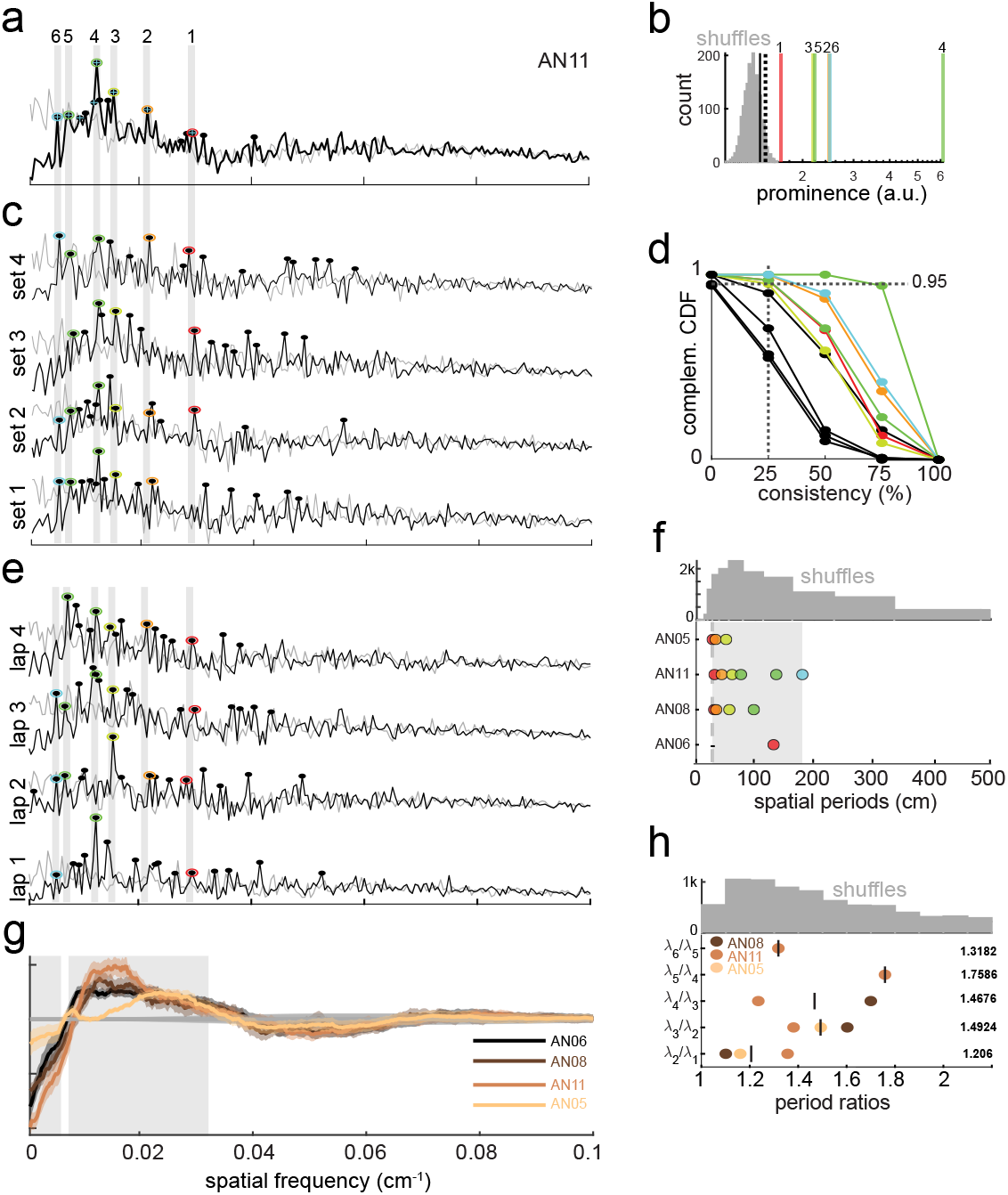
Evidence of grid-driven structure in place cell population responses. **a**, Sum of PSD of a population of place cells’ activity over all laps in mouse AN11. The gray PSDs are obtained by randomly shuffling the positions of the place fields of all cells. Black dots indicate the top 15 most prominent peaks. Among the top 10 of these peaks, only the ones that are (1) significantly more prominent than in signals with shuffled place field locations (indicated with blue crosses, see histogram in b) and (2) consistent across cell partitions (see complementary CDF in d) are selected. In AN11, this leads to 6 selected peaks that are color-coded and numbered (exact positions given in Table S1). **b**, Distribution of peak prominence in PSD of signals with shuffled place field locations. Vertical lines indicate the significance threshold (dotted black), the prominence of selected peaks in AN11 (colored and numbered) and the prominence of peaks that are not selected because their prominence is not significantly larger than in shuffles (black unnumbered). **c**, PSDs of all the cells partitioned into 4 subsets in AN11. Only the peaks that are consistent across 100 such partitions are kept, with a significance level of 5% (right complementary CDF). **d**, Complementary CDF of the consistency of the top 10 peaks across 100 cell partitions in AN11. Selected peaks (color-coded) have at least 95% probability to be consistent across these partitions. The other peaks (black) are excluded. **e**, PSD of a population of place cells’ activity for each lap in mouse AN11. **f**, Prominent spatial periods associated to PSD peaks that are significant and consistent across partitions. Top: distribution of the most prominent spatial periods in shuffled signals (with 1000 shuffles; for each animal, we select the same number of prominent peaks from each shuffle as identified for the corresponding non-shuffled result). Given the shuffle distribution, the probability of generating data localized within the shaded interval is small (*p* = 0.013). **g**, Shades of brown: Averaged smoothed PSD across laps and cells for one animal, followed by subtraction of the mean of the shuffle PSD; grey: average shuffle PSD; thus each curve represents deviations of recorded PSD from the shuffle. Each point on the curve is a moving mean of the PSD using windows of 0.01 cm^−1^, together with the moving s.e.m. (thick ligher curve). Gray boxes indicate the spatial frequencies along which the moving mean of the PSD averaged across animals, laps and cells differs from the corresponding shuffle PSD by at least two s.e.m. **h**, Ratios of successive spatial periods in all mice. The mean ratios are given on the right. Top: similar to f, distribution of ratios of successive spatial periods in shuffled signals, given which it is very unlikely to generate ratio data confined to the observed interval (*p* = 0.0005).

We next applied a more stringent condition to test the selection of peaks: we computed the PSDs by averaging across the whole population, but split results by lap runs. Since each lap is done at a different time, and neural responses are subject to independent path integration errors in each lap and other potential non-stationarity across laps, this is a more stringent criterion. We found that the selected peaks were nevertheless consistent across laps, Fig. 4e.

These analyses together yielded six peaks corresponding to spatial periods of 33.9, 45.98, 63.49, 78.43, 137.93 and 181.82 cm (Fig. 4f and Table S1), over an analyzed PSD range that could have yielded periods from 10 cm to 4000 cm (frequency range of (0.1 m)^−1^ to (40 m)^−1^). Notably, the identified periods fall in the narrow range of possible grid cell periods, which span 30-200 cm [39, 40].

Applying our analysis across animals, we found a small set of prominent, significant, and consistent peaks, all falling in the range of 30-200 cm (Fig. S5 and Table S1). To consider whether these peaks might be driven by external features in the environment or an experimental artefact, we compared the specific PSD peaks across animals to find that although peaks are within the range of grid periods, they are not precisely aligned, meaning that they cannot be attributed to spatially periodic external cues or other periodic artefacts in the experimental process (Table S1). We also computed the PSD of the external visual stimulus and the reward schedule (Fig. S6), showing that these (intentionally randomized features of the task) do not exhibit significant peaks, including specifically within the grid cell period band.

We compare the set of prominent PSD peaks with the distribution of PSD peaks expected from shuffles that preserve the number of fields per cell but not the field positions (details in Methods), Fig. 4f, and find that while the shuffle distribution has long tails, no data lie in those tails; the probability of generating the data in the observed interval from the shuffle distribution is small (*p* = 0.013). Consistent with the finding of individual PSD peaks localized within the grid cell frequency band in all animals and the model predictions (Fig. S3c), an examination of the PSD envelope obtained from averaging across all animals, laps, and cells reveals an overall under-representation of large spatial periods compared to the shuffle PSD (cluster spanning [1.67, 40] m), and an overall over-representation of an interval consistent with grid periods (cluster spanning [31.25, 129] cm), Fig. 4g. Besides, as described in [40], prominent peaks in the PSDs could also be reflected in inter-field-interval (IFI) distributions. We confirm that IFIs distributions in the experiments contain prominent, significant and consistent peaks that are consistent with the PSD results and the presence of a structured grid response, however the signal-to-noise ratio of the IFI compared to the PSD prevented us from identifying an equal number of robust peaks (Fig. S7). This is because, at the opposite of the PSD, the IFIs only account for the distances between adjacent fields, throwing away higher-order regularities.

Finally, we compute the within-animal adjacent period ratios based on the selected PSD peaks; these ratios fall between 1.1 and 1.8, consistent with the period ratios observed for adjacent grid modules [39]. The distribution of prominent peak ratios obtained from shuffles (Methods) also has long tails, Fig. 4h, with the ratio data significantly different from that predicted from the shuffled ratio distribution (*p* = 0.0005).

## Discussion

The origin of CA1 activity is a critical puzzle in our understanding of episodic memory [26]. Though we are still many steps from the ultimate goal of understanding hippocampal memory organization well enough to predict most place cell firing fields, we have unmasked a key structured component in the CA1 place cell population response that is grid-cell driven and imposes order on the seemingly random forest of fields. It was possible to uncover a structure that had previously been hidden in data because of the specific focus on recording large enough neural populations in large enough featureless environments, motivated by theoretical predictions and modeling. Notably in terms of process, for this work we first built models and generated the central predictions about PSD peaks in the hippocampus, then subsequently designed and performed the experiments and analyses specifically to test the specific predictions. This procedure mitigates the serious problems arising from multi-hypothesis testing [41] and that fact that it was possible to first make then test such a prediction highlights how Neuroscience is emerging as a firmly quantitative and theoretically-driven discipline.

The revealed CA1 scaffold can enable the learning of many distinct spatial cognitive maps by driving a large set of non-repeating population-level place field arrangements [24] that can then be associated with cues in the traversed space to represent the bindings between “what” and “where” within the space. Since grid cells also increment their states in response to traversal of non-spatial cognitive spaces, place cells can use their grid-driven scaffolds to map (and perform movement-based inference on [42]) these spaces in the same way.

Our findings suggest that grid cells drive place cells – providing a common structure to the whole CA1 population that becomes visible by averaging across place cells – as in the framework of Fig. 1a. For concreteness, we emulated connectivity and synaptic weights based on data reported in the literature, and we assumed a fixed common activation threshold across place cells. However, within the theoretical framework in which place cell responses are derived though grid and non-grid cue conjunctions, a diverse range of model variants can also reproduce random-looking place field statistics with a population-level scaffold, including those with: cellular heterogeneity, Hebbian learning in grid-to-place weights, or competitive *K*-winner-take-all lateral interactions among place cells that adaptively determine activation thresholds. The scaffold prediction is insensitive to such variations in model detail and the addition of non-grid inputs because these factors average out at the population level. In other words, it is a general result about the structure of place fields assuming some grid cell contribution, and should hold for most grid-driven place cell models [43, 44, 45, 42].

Our results contribute to an emerging body of work revealing the presence of pre-configured states in hippocampus that are subsequently harnessed for memory [28, 29, 3, 27], and suggest that these pre-configured states may be determined and structured by the grid cell input to hippocampus. Thus, instead of creating de-novo new stable memory states to store a new experience, the system may pick internal representations from an internal scaffold of largely pre-defined states. Although it is not yet understood how and to what extent these internal states are modified and updated by experience [29], our work provides concrete evidence for such pre-configured scaffolds in the CA1 population that constrain future attempts to understand the brain’s hippocampal memory system.

## Methods

### Inputs to place cell population model

In the 1D environment, we simulated *N*_*PC*_ = 253 place cells (PCs) when a virtual animal was traversing a 48m long track, as in [23]. Each PC can potentially receive inputs from grid cells (GCs), non-grid spatial cells and noise.

To simulate quasi-periodic grid responses in 1D, we first generate 2D responses with Gaussian tuning with standard deviation 0.212 *λ*_*m*_ on a hexagonal lattice, with spacing *λ*_*m*_ and orientation *θ*_*m*_ for module *m* [36]. 1D responses of GCs from the same module are then generated as parallel 1D slices of this lattice as in [36], with phases regularly distributed. In all simulations, we use three modules of *N*_*GC*_ = 1600 grid cells with grid spacings 43, 61 and 83 cm and orientations 0.39, 0.46 and 0.03 radians (leading to a total of 4800 GCs).

Non-grid spatial responses are simulated as Gaussian-profiled activity bumps at random locations on a 1D track. The widths of the bumps follow the widths of the grid cell activity bumps in the three modules. There are as many non-grid spatial cells as grid cells (4800).

Noise mimics the unknown and non-spatial sources of input to place cells; it is modeled as independently drawn Gaussian white noise with no across-lap consistency.

### Path-integration errors

To introduce path-integration errors in GCs, the perceived location along the track at step *i* is 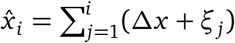. The initial perceived location is 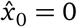 cm; displacements at each step are *Δx* = 1 cm, and the noise introduced in estimating each displacement is 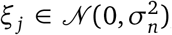, with *σ*_*n*_ = 1 cm in Fig. S3b. The true location along the track at step *i* is 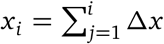 with initial location *x*_0_ = 0 cm. Thus, the resulting grid cell activity deviates by an increasing amount along the track from a perfect 1D slice through a 2D grid [36].

### Connectivity matrix in model

Connection weights are drawn from a log-normal distribution with parameters *µ* = 0 and different *σ* to adapt the weight skewness (*σ* = 1.2 to generate the results of Figs. 2 and S2a-b-c) [33]. Connection probability *c* determines the fraction of non-zero weights, which is implemented by randomly setting 1 − *c* fraction of all possible weights to zero. It is set to 0.25 to reach a realistic number of synapses from GCs to PCs given the number of GCs [46, 47]. For the spatial non-grid inputs, connectivity is set to 0.75 to account for more projections than those coming from GCs [47]. This leads to an average of 1200 GC and 3600 non GC inputs to each PC. Postsynaptic L1 normalization is imposed (all weights projecting to a PC sum to 1).

The relative contributions of the different types of inputs (grid, non-grid and noise inputs) can be varied and are quantified in terms of grid-to-non-grid and grid-to-noise ratios, expressed in 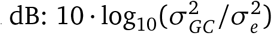, where the GC variance 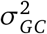 and the non-grid or noise variance 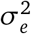 are computed over space and all simulated PCs.

### Defining place fields in model

The total input into a PC is the sum of all grid, non-grid and noise inputs. To define a place field, we set a threshold on the summed input; locations with activity above the threshold can potentially be a place field. The threshold, which we set to be the same across PCs, can be viewed as inhibition within the PC local circuit. Alternatively, an individual threshold could be drawn for different PCs, to simulate potential biophysical diversity across cells [34].

The widths of locations with activity above threshold vary. Too narrow activity above threshold is not considered as a place field, and we set a width constraint of 15 cm [23].

### Two-photon calcium imaging of mice CA1 on a VR track

Four mice (AN05, AN06, AN08, AN11) were trained to run on a 40 cm-diameter spherical treadmill at the center of a half-cylindrical screen onto which monochromatic virtual environments were projected (Fig. 3a) [34, 48, 49, 50]. The animal’s view of the environment was rendered with custom virtual reality software [50] and updated based on treadmill movement. We designed an environment with a ∼40 m-long track with randomly placed proximal cues on the walls to provide optic flow (Fig. 3b). The mice were trained for more than a month to run straight on this long track, and the environment was therefore familiar to them. There was a tunnel-like structure at the start of the track to indicate the beginning of the track for each lap. Behavior consisted of running the entire track’s length in one direction then being teleported to the start for another lap, for a total of 4 laps in the given environment (mouse AN05 did 9 laps). The proximal cues were black stripes on a white background with randomly sampled tilts of 25, 35 or 45°. The stripe widths were uniformly distributed between 2 and 20 cm and followed by a white interval of the same width, identical for all laps in all four mice. Identical distal cues (four mountains) were presented in all laps to mice AN08 and AN11, but no distal cues for mice AN05 and AN06.

We had to make the mice run straight and in the middle of the track during the experiments. To do so, there were visible baffles of 5 cm every 40 cm for AN05, the first trained mouse. This regularity was accounted for in the analyses (see below). To avoid using such periodic visual stimuli which induce cue-driven periodic neural responses, all visible baffles were removed for the later mice, and there were instead invisible slow lanes on both sides of the track (see Fig. 3b) for AN06, AN08 and AN11. Water reward was provided at random locations that changed for each lap. In the first experiment with mouse AN05, possible reward spots were evenly spaced by 20 cm and reward was delivered randomly in 1 out of every 10 spots. In order to avoid any partial regularity in reward locations, the reward locations were further randomized for the next mice. For mice AN06, AN08 and AN11, reward was delivered with probability 10% every 5 cm. Despite the reward regularity and the presence of periodic visual cues for AN05, this first mouse was kept for the analyses as it lead to similar conclusions. We imaged the calcium activity of dorsal hippocampal CA1 pyramidal neurons in transgenic mice (*n* = 4) expressing GCaMP6f (GCaMP6f-expressing cells do not express GAD67) with a sampling rate of 30 Hz. In mice AN06, AN08, AN11 and AN05, respectively 630, 670, 550 and 650 neurons were identified in each imaging session.

The raw fluorescence calcium signals of each cell were denoised by (1) applying a third-order Savitzky-Golay filter and (2) thresholding the obtained traces to only keep the calcium transients higher than the sum of the median value and three interquartile ranges of activity within sliding windows of ± 8 seconds around each time point [51].

### Defining place fields in data

The fluorescence transients as a function of time are first interpolated and expressed as a function of the spatial position along the track with 1 cm resolution. Then, transients observed within 15 cm windows of each others are grouped, and a place field is defined with a minimum-width constraint of 5 cm. We consider each significant transient of sufficient size to be a place field without accounting for repeatability across laps, because the long and locally featureless track is expected to induce drifts (PI errors) in spatial estimation that can vary across laps.

### Place field structure analysis in both data and model

In the simulation, we obtain thresholded activity for each place cell as place cell activity as described above. In both data and model, we detect the center of mass (COM) of activity that satisfies the place field constraints, setting COM location as 1 and 0 otherwise. The binary signals are convolved with Gaussian windows of 4.7 cm FWHM, yielding standardized place fields. We analyze the spatial structure of these place fields by computing the combined power spectral density (PSD) of place field distribution in a population of place cells. The combined PSD is obtained by computing the PSD of place field distribution for each neuron (and each lap for the data) and then summing them up. It is assumed that the grid pattern is fixed in an animal but differs across animals, so we do not expect to see the same structure in all mice.

To assess the spatial structure of place fields, the PSDs of neural activity are compared to PSDs of signals with shuffled place field locations. Shuffled signals are obtained by randomly placing the fields of each (real or simulated) neuron along the track, thus preserving the distribution of number of place fields per cell.

### Peak locations in PSDs: Simulations

In simulations, the grid-induced spatial frequencies are known (Fig. 1c). Indeed, the PSD of an idealized (infinite) 1D slice through a 2D grid lattice with spacing *λ*_*m*_ and orientation *θ*_*m*_ comprises 3 Fourier components at spatial frequencies 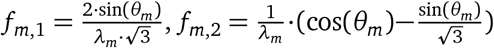 and *f*_*m*,3_ = *f*_*m*,1_ + *f*_*m*,2_ [36]. All co-modular GC responses consist of these three primary Fourier components. By linearity of the Fourier transform, the activity of an idealized place cell receiving inputs from *M* grid modules should hence contain 3*M* primary Fourier components at frequencies 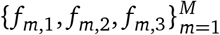. These three frequencies per module are reported with different markers in Figs. 1c-d, 2b-c and S2b-c. To quantify the extent to which these frequencies are enhanced by the GC inputs in the PC activity, we compute the discriminability index between the shuffled and real distributions of amplitudes at these frequencies as |*µ*_*r*_ −*µ*_*s*_| */*(0.5·(*σ*_*r*_ + *σ*_*s*_)), where the subscripts *r* and *s* refer to the real simulated and shuffled signals respectively [52].

### Peak locations in PSDs: Analytics

We expect to observe GC-driven frequencies as prominent peaks in the activity of PCs if the PSDs of a large number of PCs are summed, as we show next. We consider that the firing rate of a PC *µ* in lap *l* is given by 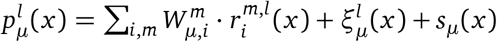, where 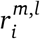 is the firing rate of GC *i* in module *m* in lap *l*, 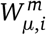 are random projection weights (≥ 0, with mean *µ*_*W*_ and variance 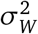), *s*_*µ*_(*x*) represents non-periodic (non-grid) but spatially anchored and reliable inputs (thus no lap-dependence), and 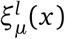 is the spatially unreliable noise (i.e., assumed to vary from lap to lap). Using the symbol ℱ to denote a Fourier transform, the PSD of a PC in one lap is given by:

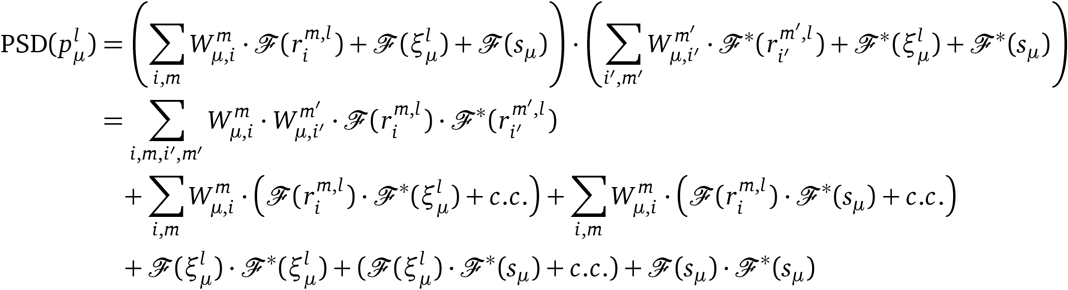

We compute the expectation value ⟨⟩_*W*_ over the random weights *W*, to obtain:

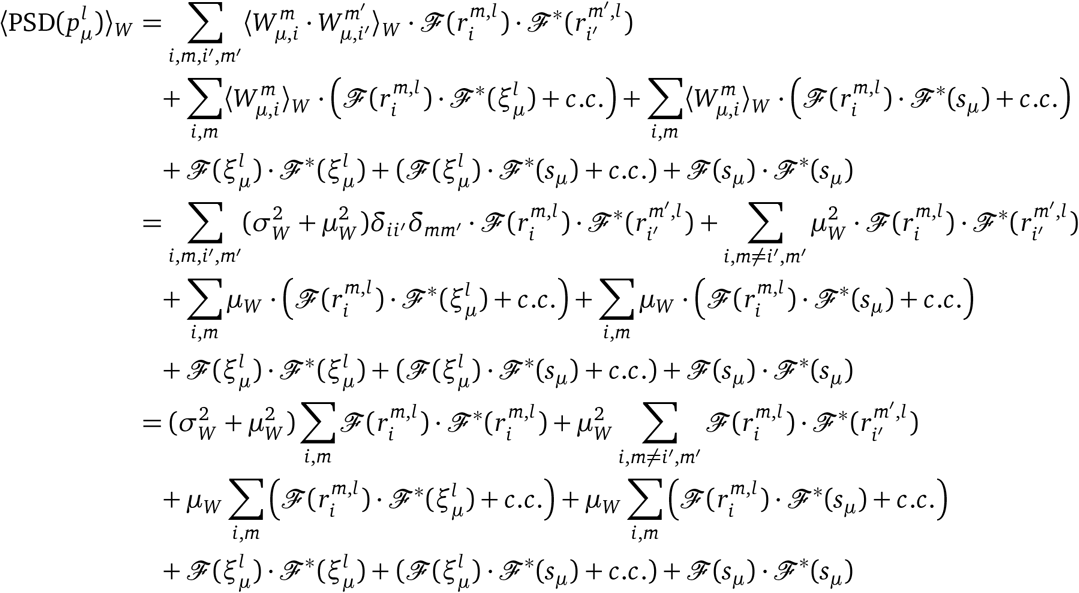

Note that 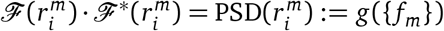 which is independent of *i* (cell index) across all cells within a module [40]. In addition, the Fourier transform 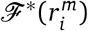 is highly sparse, consisting of only a few non-zero peaks at frequencies {*f*_*m*_} ≡ {*f*_*m*,1_, *f*_*m*,2_, *f*_*m*,1_ + *f*_*m*,2_} as described above, that are determined by the periodicity *λ*_*m*_ and slice orientation *θm* of the module [40], so we also have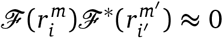 for *m ≠m′*. Finally, note that the Fourier transform of a white noise term is also a white noise, thus 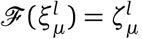, where *ζ* is a white noise term with the same variance as *ξ*. With these three facts, we have:

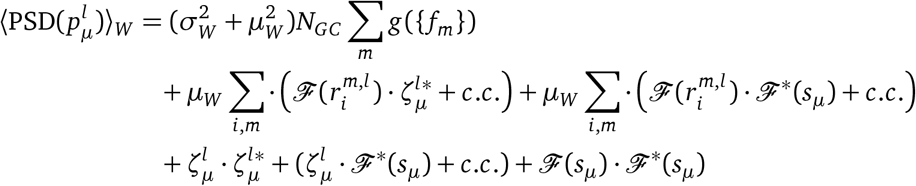

We now compute the sum over all place cells and laps:

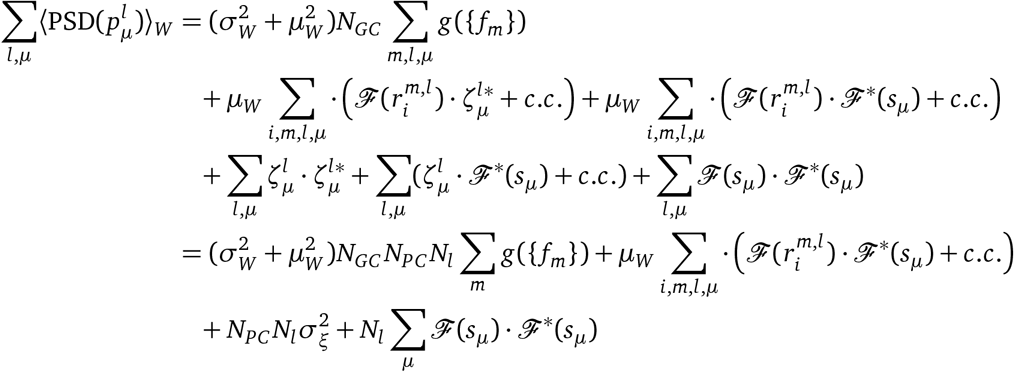

where for the second equality we have used the fact that 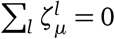. Note that for any given place cell *µ*, the spatial non-grid inputs are relatively sparse and randomly arranged compared to grid inputs, and thus the Fourier power spectrum ℱ (*s*_*µ*_) ℱ^*^(*s*_*µ*_) = *g*({*f*_*µ*_}) will consist of relatively broad features in Fourier space. Though broad, the Fourier features are not merely a DC component for a single cell. However, assuming that the non-grid spatial inputs are relatively uncorrelated across the population of cells, the sum of the power spectral density of these inputs over a large number of place cells will be approximately constant, 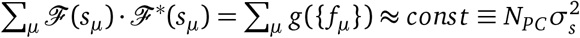, where 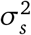 is defined as the power of the non-grid spatial inputs to each place cell. Finally, note that in the absence of path integration errors, the terms 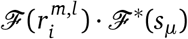 pick out the elements of the broad ℱ^*^(*s*_*µ*_) that are aligned with the Fourier frequencies of the grid cells, while path integration errors induce phase shifts that can smear out the selection of the grid peaks. In total, this term is smaller than the first term on the right hand side of the above. Thus, we have:

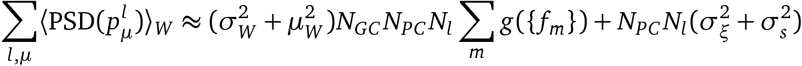

Hence, in the sum across place cells and laps, the term containing the grid cell PSDs (*g*({*f*_*m*_})) adds coherently and with structured Fourier peaks, while the rest of the terms contain little systematically occurring frequency content and wash out to a total contribution that amounts to an overall DC offset in Fourier space. The grid cell Fourier peaks also sum constructively across all grid cells within a module that project to each place cell (*N*_*GC*_), in addition to summing across place cells (*N*_*PC*_) and laps (*N*_*l*_). Since *N*_*GC*_≫ 1 (probably numbering in the tens to hundreds), it follows that in the sum there should be prominent PSD peaks that correspond to the grid module spatial frequencies.

### Peak locations in PSDs: Data analysis

Topographical prominence can be used to characterize and locate the peak locations in PSDs. In experimental data, we select the top 15 most prominent peaks in each lap and animal. In AN05, the first trained animal, the frequencies corresponding to the periodicity of the visible baffles positioned every 40 cm are excluded from this selection (0.025 cm^−1^ and harmonic frequencies). We then only keep the peaks that are among the top 10 peak locations in the averaged PSD per mouse and that are (1) consistent across 100 random partitions of the cells into 4 subsets (minimum consistency of 50% and tolerance of 3 frequency bins or 0.00075 cm^−1^) and (2) significantly more prominent compared to distributions of prominence in signals with shuffled place field locations (*n* = 2000 random placements of the fields for each cell, lap and mouse, using the same constraint for the minimum distance between adjacent fields as for the real data). Significance level is set to 5% and is adapted for multiple comparisons with the Holm-Bonferroni correction for each animal. To improve visualization, the PSDs are baseline-corrected in the figures, after the peaks are selected and assessed (so that it only improves the readability of Fig. 4 without affecting the results in any way).

Finally, to ensure that the identified prominent frequencies are not driven by environmental features, we look for regularities in the visual cues and reward locations that could have induced artifactual prominent PSD peaks in neural activity (see Fig. S6). We check whether any prominent spatial period is found both in neural activity and in visuals or rewards (with a tolerance of 3 frequency bins for matching). For the visuals, we use screen shots of the track as seen by the mice every 1 cm, compute the spatial autocorrelation of these screen shots and then the PSD of this autocorrelation. The top 15 most prominent peaks in this PSD are selected. For the reward locations, we compute, for each animal and each lap, the PSD of the binary reward signal, which is set to 1 when there is a reward and 0 otherwise, with a 1 cm spatial resolution. The top 15 most prominent peaks are selected in the PSD for each lap and in the PSD averaged over the laps. Then, we keep the peaks that are among the top 10 peak locations in the averaged PSD per mouse and that are (1) consistent across the 4 laps (minimum consistency of 50% and tolerance of 3 frequency bins or 0.00075 cm^−1^) and (2) significantly more prominent compared to distributions of prominence in binary signals with shuffled reward locations (*n* = 2000 random placements for each lap and mouse). After comparing the selected peaks in neural data, rewards and visuals, we solely excluded one prominent peak at 266 cm in AN05 which was selected in both neural activity and visuals. No other matching was found.

### PSD envelope in data

In addition to assessing the precise locations at which prominent peaks are observed in the recordings, we test whether the envelope of the experimental PSD differs from the PSDs of shuffled signals, as expected when averaging PSD with variable PI errors (Fig. S3c). To this end, we compute the average PSD across animals, laps and cells and compare this curve to a matched shuffled PSD averaged over animals, laps, cells and 1000 repetitions of the field placements for each recording. These two curves are then compared using a moving mean and moving s.e.m. with a 0.01 cm^−1^ frequency window.

### Peak locations in inter-field-interval distributions

In addition to the PSD analysis, we check whether or not we can identify prominent spatial periods by looking at the distributions of inter-field-intervals (IFIs). If an idealized place cell combines grid inputs from different modules *m*, IFIs of 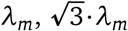 and 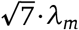 are expected to be overrepresented [36]. The distribution of IFIs are computed across the cells for each lap individually and by merging all laps using a bin width of 1 cm. For each histogram, the top 15 most prominent peaks are identified. Among these peaks, we select the significant and consistent ones in the same way as for the PSD peaks (Fig. S7). Although the selected peaks fall within a range which is consistent with a grid drive, the signal-to-noise ratio of the IFI compared to the PSD prevents from robustly identifying as many peaks as in the PSDs.

## SUPPLEMENTAL INFORMATION

Supplemental Information includes 7 figures and 1 table and can be found with this article online.

## ACKNOWLEDGMENTS

This work was supported by the Simons Foundation through the Simons Collaboration on the Global Brain, the ONR, the Howard Hughes Medical Institute through the Faculty Scholars Program to I.R.F., and the Alfred P. Sloan Research Fellowship FG-2017-9554 to T.T.. M.Y.Y. thanks Ernie Hwaun and Jens-Oliver Muthmann for discussions.

## AUTHOR CONTRIBUTIONS

Conceptualization, D.M., M.Y.Y., T.T., and I.R.F.; Experiments: J.S.L and A.K.L.; Data Analysis & Model Simulation, D.M., M.Y.Y; Writing, D.M. and I.R.F. with input from all co-authors.

---

**Fig. S1:**
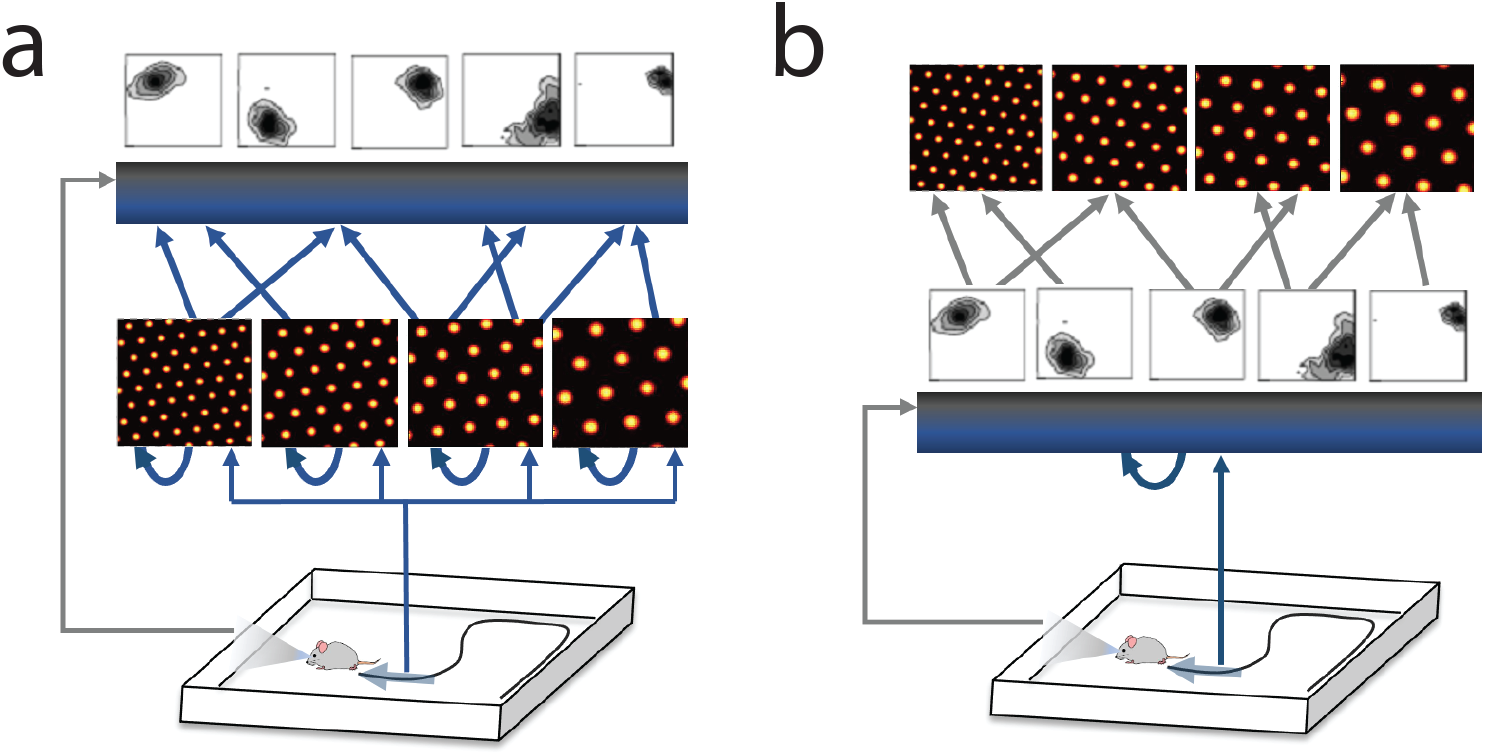
Models linking activity in grid cells and place cells. During spatial navigation, place cells receive sensory inputs (vertical gradient) and can be modeled to primarily either combine these external inputs with internal grid cell inputs (**a**), or rather drive the grid cells activity (**b**).

**Table S1:**
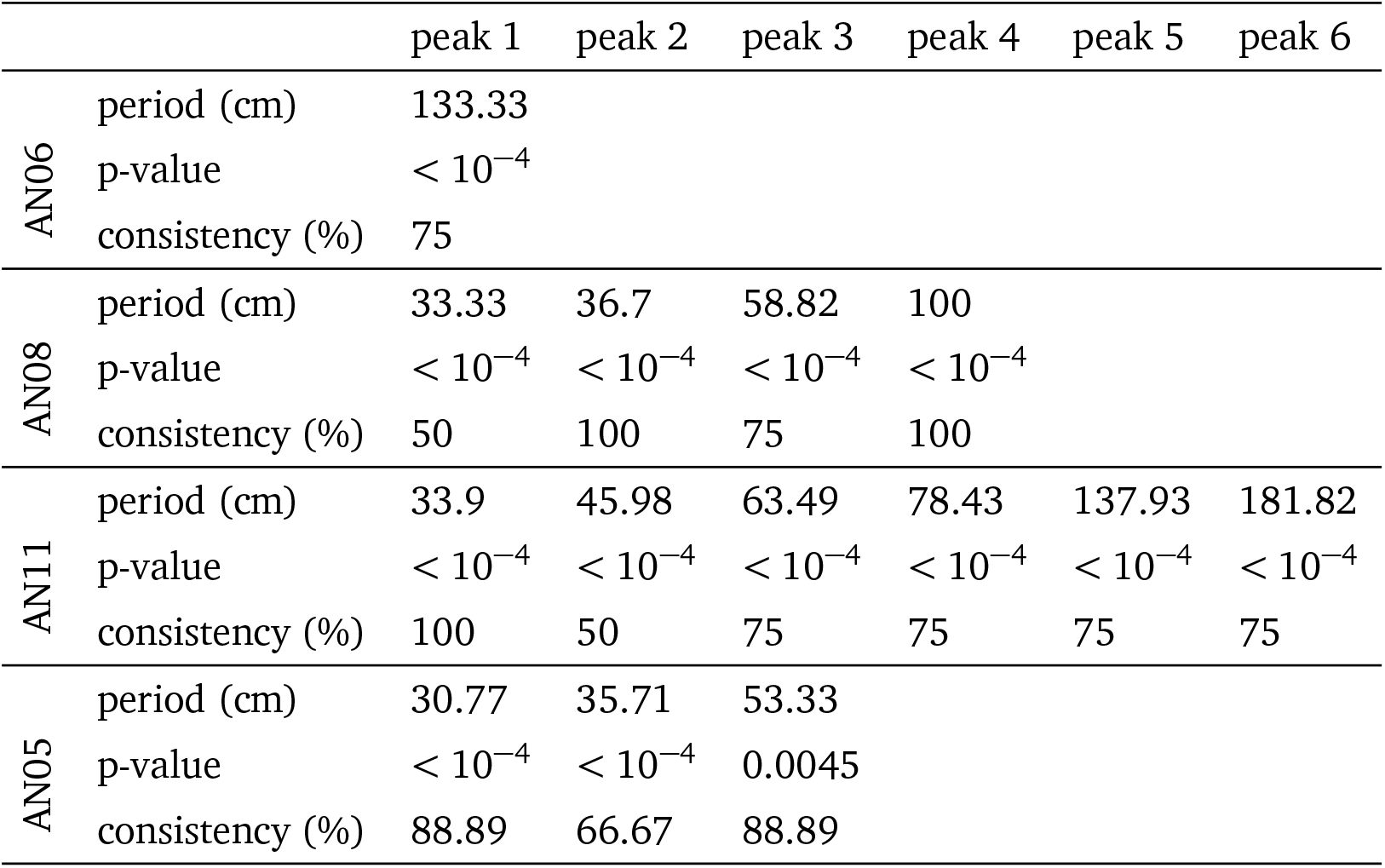
Prominent spatial periods in place cell activities. Spatial periods associated to PSD peaks that are significant and consistent across cell partitions. Significance is computed based on distributions of the most prominent peaks in signals with randomized place field locations (2000 random placements). Consistency is the percentage of laps in which the prominent peaks are observed.

|  |  | peak 1 | peak 2 | peak 3 | peak 4 | peak 5 | peak 6 |
| --- | --- | --- | --- | --- | --- | --- | --- |
| AN06 | period (cm) | 133.33 |  |  |  |  |  |
| | p-value | $< 10^{-4}$ | | | | | |
|  | consistency (%) | 75 |  |  |  |  |  |
| AN08 | period (cm) | 33.33 | 36.7 | 58.82 | 100 |  |  |
| | p-value | $< 10^{-4}$ | $< 10^{-4}$ | $< 10^{-4}$ | $< 10^{-4}$ | | |
|  | consistency (%) | 50 | 100 | 75 | 100 |  |  |
| AN11 | period (cm) | 33.9 | 45.98 | 63.49 | 78.43 | 137.93 | 181.82 |
| | p-value | $< 10^{-4}$ | $< 10^{-4}$ | $< 10^{-4}$ | $< 10^{-4}$ | $< 10^{-4}$ | $< 10^{-4}$ |
|  | consistency (%) | 100 | 50 | 75 | 75 | 75 | 75 |
| AN05 | period (cm) | 30.77 | 35.71 | 53.33 |  |  |  |
| | p-value | $< 10^{-4}$ | $< 10^{-4}$ | 0.0045 | | | |
|  | consistency (%) | 88.89 | 66.67 | 88.89 |  |  |  |

**Fig. S2:**
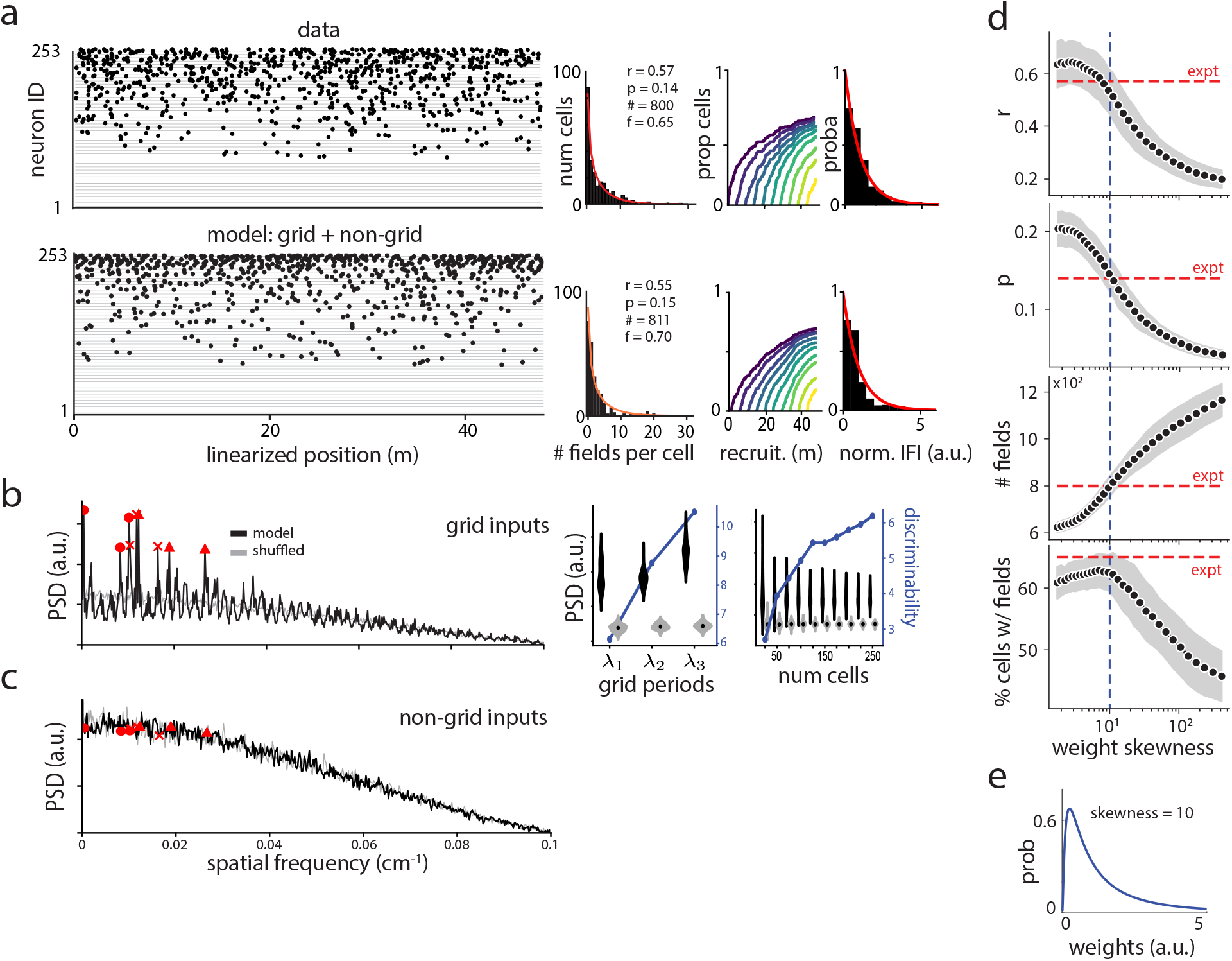
Field statistics and structure in activity simulated by the model. **a**, Place fields of populations of 253 place cells observed in electrophysiological recordings (row 1, as reported in [23]) and generated by our model with grid cell and non-grid spatial cell inputs (row 2). The distributions of the number of fields per place cell are well approximated by Gamma-Poisson distributions (red curves). r, p: Gamma-Poisson shape parameters, #: total number of place fields, f: fraction of cells having at least one field along the track (%). The recruitment of place cells along the track is memoryless, and normalized inter-field-intervals follow an exponential distribution (red curve fitted on cells with at least 6 fields among 253 simulated cells). **b**, Similar to Fig. 2, PSD of place field distribution with GC inputs alone. Violin plots indicate distributions of PSD amplitudes at GC-related frequencies with *n* = 30 simulations. **c**, PSD of place field distribution with only spatial non-GC inputs, in 253 place cells. **d**, Field statistics as a function of the skewness of the lognormal distribution of the connection weights from GCs to place cells, with GC inputs only. Horizontal dotted lines indicate the parameters values (Gamma-Poison parameters r and p, total number of fields and fraction of cells with at least one field) observed in empirical recordings from [23]. Vertical lines pinpoint the fixed skewness which enables reaching all these parameters. Intervals of one standard deviation around the mean are shown for 30 random samplings of the weights. **e**, Distribution of the weights with skewness 10 which leads to realistic field statistics.

**Fig. S3:**
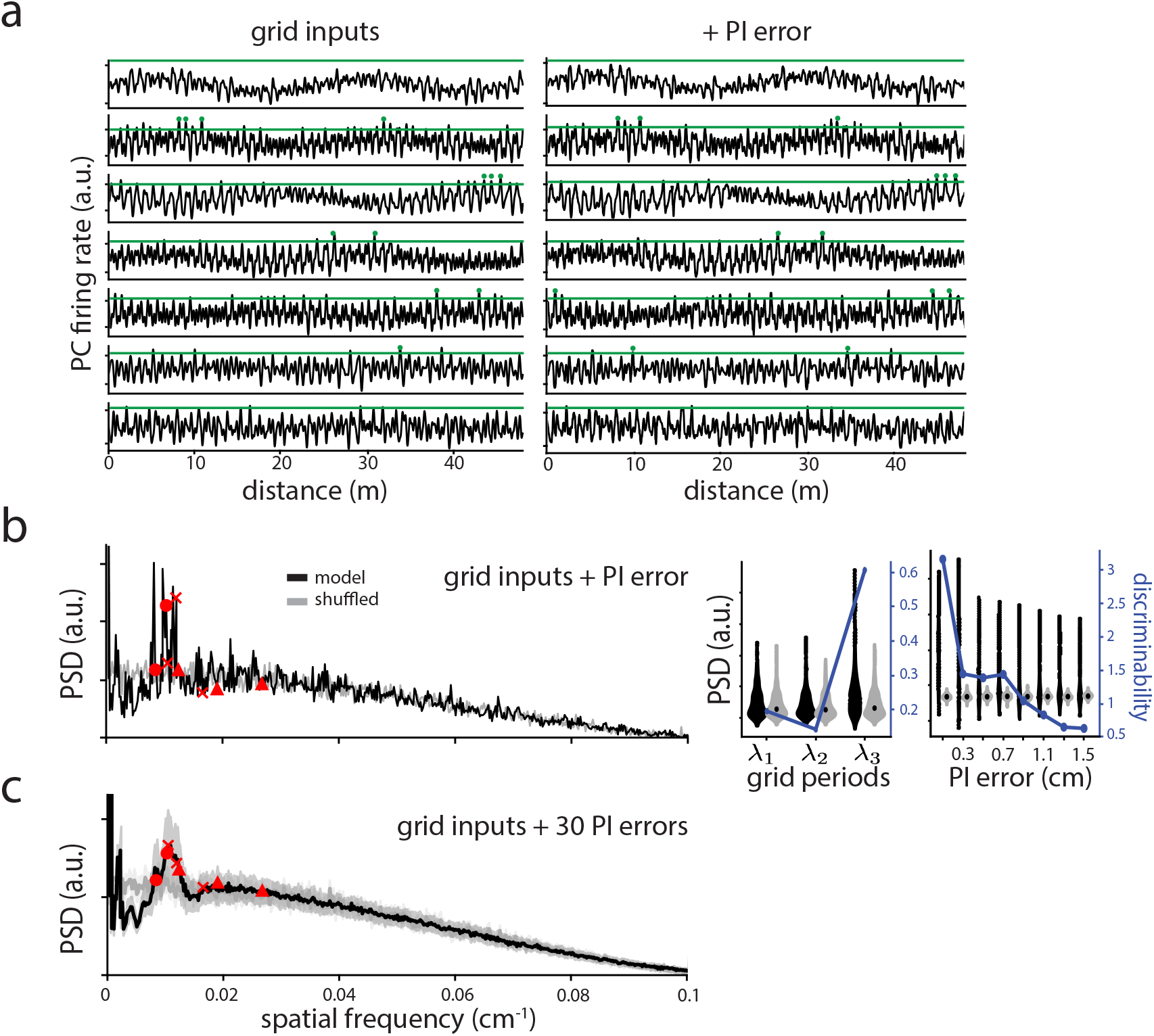
Effects of path-integration (PI) errors on predicted place field structure. **a**, Firing rates of example place cells along the track with GC inputs, without (left) and with (right) path-integration (PI) errors with 1 cm standard deviation. Accumulating PI errors are expressed in terms of s.d. per cm of displacement along the track (see Methods). A threshold (green line) is applied to define place fields (green dots). **b**, Similar to Fig. 2, PSD of place field distribution with GC inputs and cumulative PI errors with 1 cm standard deviation, in 253 place cells. Even at very large PI values (standard deviation of 1 or 1.5 cm per cm of displacement), some GC-induced peaks remain discriminable at the population-level, as indicated by the prominent peaks in the PSD (left) and the extent of the violin plots (right). **c**, Similar to b, with an average over 30 simulations of PI errors with 1.5 cm s.d. (shaded areas give the s.d.). The different PI errors lead to a PSD envelope which deviates from the shuffle PSD with a hump over GC-related frequencies and an under-representation of smaller spatial frequencies. The main hump is expected to get broader if PSDs from place cells driven by different sets of grid modules (e.g. in different mice) are averaged.

**Fig. S4:**
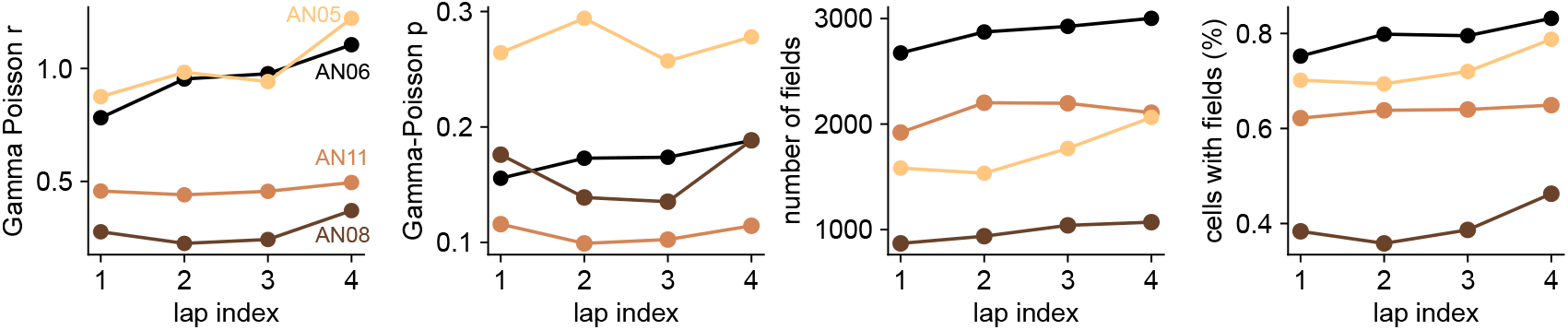
Place field statistics across laps. Features of the place field distributions for all 4 mice across 4 laps along the VR linear track. The place field statistics are mostly stable across laps per animal, with a slight tendency towards an increased number of place fields with the lap index.

**Fig. S5:**
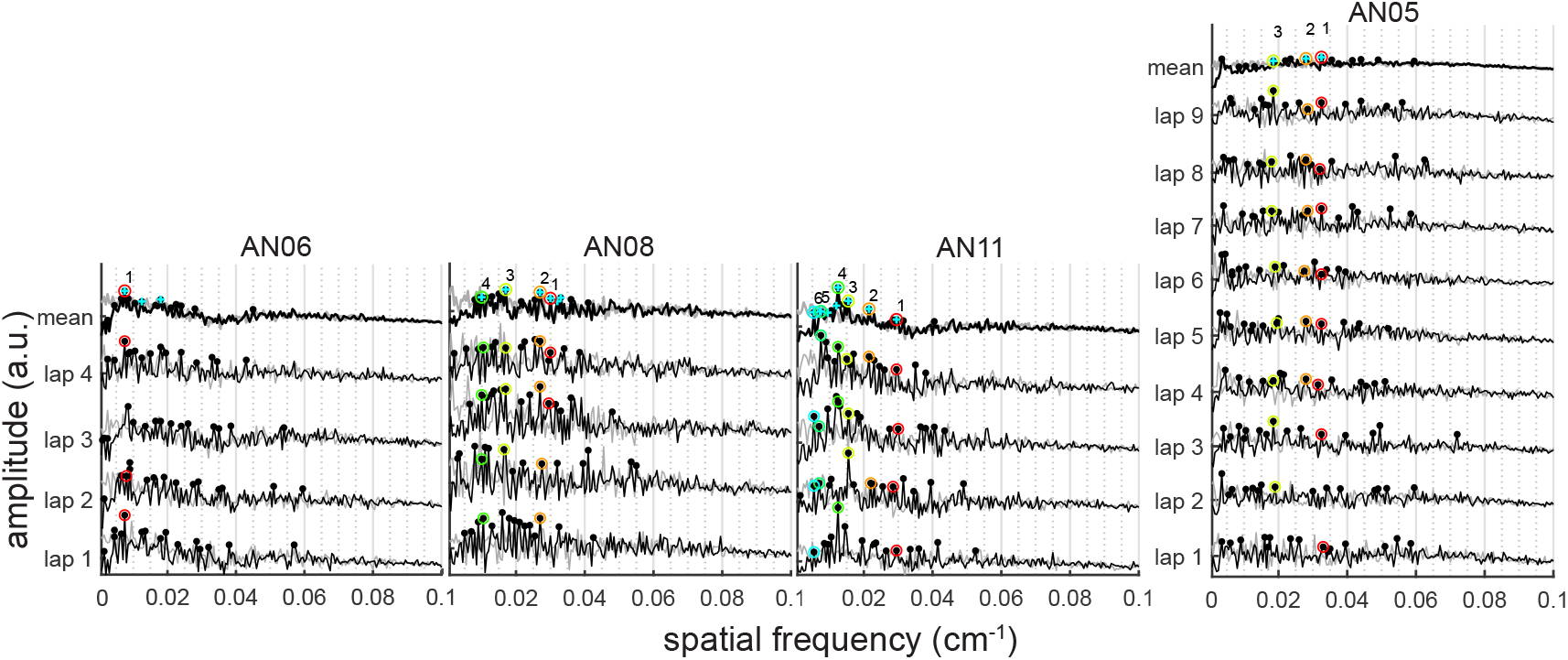
Grid-driven structure in place cell population responses. PSD of populations of place cells’ activity across the laps in all mice. Black dots indicate the top 15 most prominent peaks. Among these peaks, only the ones that are significantly more prominent than in shuffled signals (as indicated with blue crosses) and consistent across cell partitions are selected (color-coded and numbered peaks, see Table S1).

**Fig. S6:**
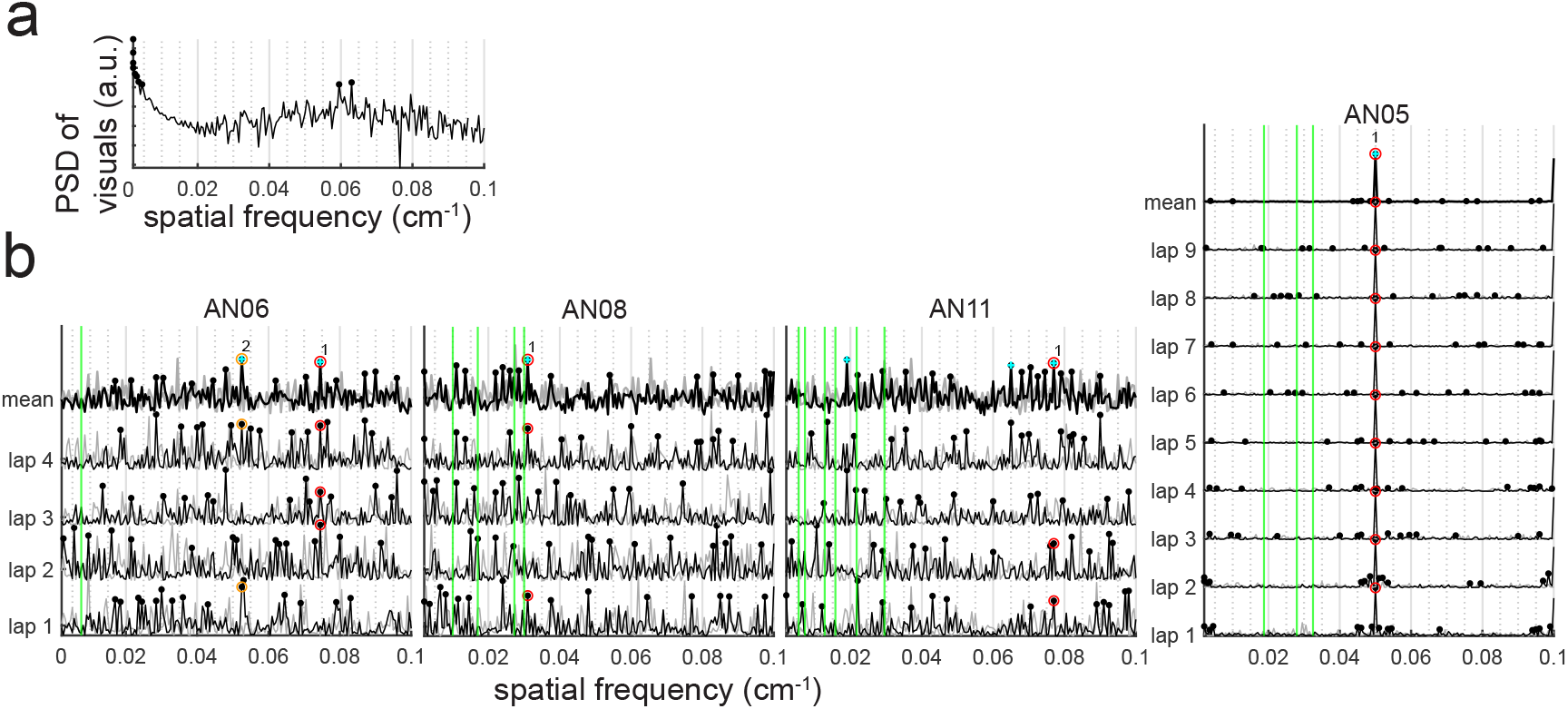
Structure in place cell population responses is not induced by rewards or visual cues. **a**, PSD of the visuals along the track. Black dots indicate the top 15 most prominent peaks. These prominent peaks are located at spatial periods that are very different from the ones picked in PSDs of neural data (Fig. S5). **b**, PSD of reward locations across the laps in all mice. Black dots indicate the top 15 most prominent peaks. Among these peaks, only the ones that are significantly more prominent than in shuffled signals (as indicated with blue crosses) and consistent across the laps are selected (color-coded and numbered peaks). The vertical lines show the positions of the selected prominent peaks in the PSD of neural activity from Fig. S5. These prominent spatial frequencies are not associated with regularities in reward locations. AN05, the first mouse to be trained, had partially non-random reward locations spaced every 20 cm, which explain the large peak at 0.05 cm^−1^ in the last PSD (see Methods).

**Fig. S7:**
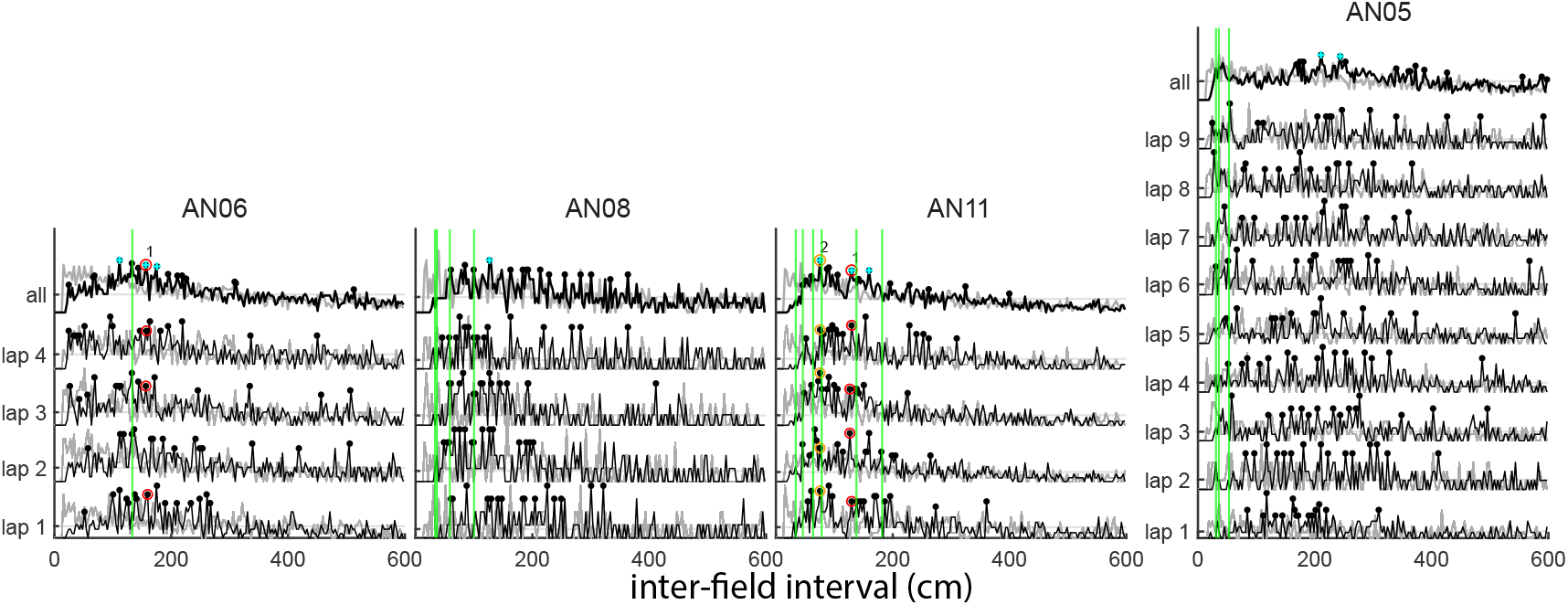
Grid-driven structure in place cell population responses based on inter-field-intervals. Inter-field-intervals (IFIs) distributions in populations of place cells’ activity across the laps in all mice. Black dots indicate the top 15 most prominent peaks. Among these peaks, only the ones that are significantly more prominent than in shuffled signals (as indicated with blue crosses) and consistent across cell partitions are selected (color-coded and numbered peaks). The vertical lines show the positions of the selected prominent peaks in the PSD from Fig. S5.

## References

[1] Tulving, E. & Markowitsch, H. J. Episodic and declarative memory: role of the hippocampus. Hippocampus 8, 198–204 (1998).

[2] Grosmark, A. D. & Buzsáki, G. Diversity in neural firing dynamics supports both rigid and learned hippocampal sequences. Science 351, 1440–1443 (2016).

[3] Farooq, U. & Dragoi, G. Emergence of preconfigured and plastic time-compressed sequences in early postnatal development. Science 363, 168–173 (2019).

[4] Squire, L. R. Memory and brain systems: 1969–2009. Journal of Neuroscience 29, 12711– 12716 (2009).

[5] Eichenbaum, H. What versus where: Non-spatial aspects of memory representation by the hippocampus. Curr Top Behav Neurosci 37, 101–117 (2018).

[6] Wiener, S., Paul, C. & Eichenbaum, H. Spatial and behavioral correlates of hippocampal neuronal activity. Journal of Neuroscience 9, 2737–2763 (1989).

[7] O’Keefe, J. & Nadel, L. The hippocampus as a cognitive map (Oxford: Clarendon Press, 1978).

[8] Shapiro, M. L., Tanila, H. & Eichenbaum, H. Cues that hippocampal place cells encode: dynamic and hierarchical representation of local and distal stimuli. Hippocampus 7, 624– 642 (1997).

[9] Geiller, T., Fattahi, M., Choi, J.-S. & Royer, S. Place cells are more strongly tied to landmarks in deep than in superficial ca1. Nature communications 8, 1–11 (2017).

[10] Pastalkova, E., Itskov, V., Amarasingham, A. & Buzsáki, G. Internally generated cell assembly sequences in the rat hippocampus. Science 321, 1322–1327 (2008).

[11] Kraus, B. J., Robinson II, R. J., White, J. A., Eichenbaum, H. & Hasselmo, M. E. Hippocampal “time cells”: time versus path integration. Neuron 78, 1090–1101 (2013).

[12] Zhang, K. Representation of spatial orientation by the intrinsic dynamics of the head-direction cell ensemble: a theory. J Neurosci 15, 2112–2126 (1996).

[13] Xie, X., Hahnloser, R. H. R. & Seung, H. S. Double-ring network model of the head-direction system. Phys Rev E Stat Nonlin Soft Matter Phys 66, 041902 (2002).

[14] Burak, Y. & Fiete, I. R. Accurate path integration in continuous attractor network models of grid cells. PLoS Comput Biol 5, e1000291 (2009).

[15] Taube, J. S., Muller, R. U. & Ranck Jr, J. B. Head-direction cells recorded from the postsubiculum in freely moving rats. I. Description and quantitative analysis. J Neurosci 10, 420–435 (1990).

[16] Chaudhuri, R., Gerçek, B., Pandey, B., Peyrache, A. & Fiete, I. The intrinsic attractor manifold and population dynamics of a canonical cognitive circuit across waking and sleep. Nat Neurosci 22, 1512–1520 (2019).

[17] Hulse, B. K. & Jayaraman, V. Mechanisms underlying the neural computation of head direction. Annu Rev Neurosci 43, 31–54 (2020).

[18] Yoon, K., Buice, M., Barry, R., C. and Hayman, Burgess, N. & Fiete, I. Specific evidence of low-dimensional continuous attractor dynamics in grid cells. Nat Neurosci 16, 1077–84 (2013).

[19] Trettel, S., Trimper, J., Hwaun, E., Fiete, I. & Colgin, L. Grid cell co-activity patterns during sleep reflect spatial overlap of grid fields during active behaviors. Nat Neurosci 22, 609–617 (2019).

[20] Gardner, R. J., Lu, L., Wernle, T., Moser, M.-B. & Moser, E. I. Correlation structure of grid cells is preserved during sleep. Nat Neurosci 22, 598–608 (2019).

[21] O’Keefe, J. & Dostrovsky, J. The hippocampus as a spatial map. Preliminary evidence from unit activity in the freely-moving rat. Brain research 34, 171–5 (1971). URL http://www.ncbi.nlm.nih.gov/pubmed/5124915.

[22] Fenton, A. A. et al. Unmasking the CA1 Ensemble Place Code by Exposures to Small and Large Environments: More Place Cells and Multiple, Irregularly Arranged, and Expanded Place Fields in the Larger Space. The Journal of Neuroscience 28, 11250 (2008). URL http://www.jneurosci.org/content/28/44/11250.abstract.

[23] Rich, P. D., Liaw, H.-P. & Lee, A. K. Large environments reveal the statistical structure governing hippocampal representations. Science 345 (2014). URL http://science.sciencemag.org/content/345/6198/814/tab-pdf.

[24] Yim, M. Y., Sadun, L. A., Fiete, I. R. & Taillefumier, T. Place-cell capacity and volatility with grid-like inputs. Elife 10, e62702 (2021).

[25] Bittner, K. C., Milstein, A. D., Grienberger, C., Romani, S. & Magee, J. C. Behavioral time scale synaptic plasticity underlies CA1 place fields. Science 357, 1033–1036 (2017).

[26] Dragoi, G. & Tonegawa, S. Selection of preconfigured cell assemblies for representation of novel spatial experiences. Philos Trans R Soc Lond B Biol Sci 369, 20120522 (2014).

[27] McKenzie, S. et al. Preexisting hippocampal network dynamics constrain optogenetically induced place fields. Neuron 109, 1040–1054 (2021).

[28] Dragoi, G. & Tonegawa, S. Preplay of future place cell sequences by hippocampal cellular assemblies. Nature 469, 397–401 (2011).

[29] Liu, K., Sibille, J. & Dragoi, G. Preconfigured patterns are the primary driver of offline multi-neuronal sequence replay. Hippocampus 29, 275–283 (2019).

[30] Kropff, E. & Treves, A. The emergence of grid cells: intelligent design or just adaptation? Hippocampus 18 (2008).

[31] Dordek, Y., Soudry, D., Meir, R. & Derdikman, D. Extracting grid cell characteristics from place cell inputs using non-negative principal component analysis. Elife 5, e10094 (2016).

[32] Stachenfeld, K. L., Botvinick, M. M. & Gershman, S. J. The hippocampus as a predictive map. Nat Neurosci 20, 1643–1653 (2017).

[33] Buzsáki, G. & Mizuseki, K. The log-dynamic brain: how skewed distributions affect network operations. Nature Reviews Neuroscience 15, 264–278 (2014). URL https://doi.org/10.1038/nrn3687.

[34] Lee, J. S., Briguglio, J. J., Cohen, J. D., Romani, S. & Lee, A. K. The statistical structure of the hippocampal code for space as a function of time, context, and value. Cell 183, 620–635 (2020).

[35] Epsztein, J., Brecht, M. & Lee, A. K. Intracellular determinants of hippocampal CA1 place and silent cell activity in a novel environment. Neuron 70, 109–120 (2011).

[36] Yoon, K., Lewallen, S., Kinkhabwala, A. A., Tank, D. W. & Fiete, I. R. Grid Cell Responses in 1D Environments Assessed as Slices through a 2D Lattice. Neuron 89, 1086–1099 (2016). URL https://doi.org/10.1016/j.neuron.2016.01.039.

[37] Harland, B., Contreras, M., Souder, M. & Fellous, J.-M. Dorsal CA1 hippocampal place cells form a multi-scale representation of megaspace 31, 2178–2190.e6 (2021). URL https://doi.org/10.1016%2Fj.cub.2021.03.003.

[38] Gauthier, J. L. & Tank, D. W. A dedicated population for reward coding in the hippocampus. Neuron 99, 179–193 (2018).

[39] Stensola, H. et al. The entorhinal grid map is discretized. Nature 492, 72–78 (2012). URL http://www.nature.com/doifinder/10.1038/nature11649.

[40] Yoon, K., Lewallen, S., Kinkhabwala, A., Tank, D. & Fiete, I. Grid cell responses in 1d environments assessed as slices through a 2d lattice. Neuron 89, 1086–99 (2016).

[41] Nosek, B. A., Ebersole, C. R., DeHaven, A. C. & Mellor, D. T. The preregistration revolution. Proc Natl Acad Sci U S A 115, 2600–2606 (2018).

[42] Whittington, J. C. R. et al. The tolman-eichenbaum machine: Unifying space and relational memory through generalization in the hippocampal formation. Cell 183, 1249–1263.e23 (2020).

[43] Solstad, T., Moser, E. I. & Einevoll, G. T. From grid cells to place cells: a mathematical model. Hippocampus 16, 1026–1031 (2006).

[44] Sreenivasan, S. & Fiete, I. Grid cells generate an analog error-correcting code for singularly precise neural computation. Nat Neurosci 14, 1330–7 (2011).

[45] Neher, T., Azizi, A. H. & Cheng, S. From grid cells to place cells with realistic field sizes. PLoS One 12, e0181618 (2017).

[46] Megıas, M., Emri, Z., Freund, T. & Gulyas, A. Total number and distribution of inhibitory and excitatory synapses on hippocampal CA1 pyramidal cells. Neuroscience 102, 527–540 (2001).

[47] Diehl, G. W., Hon, O. J., Leutgeb, S. & Leutgeb, J. K. Grid and nongrid cells in medial entorhinal cortex represent spatial location and environmental features with complementary coding schemes. Neuron 94, 83–92 (2017).

[48] Harvey, C. D., Collman, F., Dombeck, D. A. & Tank, D. W. Intracellular dynamics of hippocampal place cells during virtual navigation. Nature 461, 941–946 (2009).

[49] Dombeck, D. A., Harvey, C. D., Tian, L., Looger, L. L. & Tank, D. W. Functional imaging of hippocampal place cells at cellular resolution during virtual navigation. Nature neuroscience 13, 1433–1440 (2010).

[50] Cohen, J. D., Bolstad, M. & Lee, A. K. Experience-dependent shaping of hippocampal CA1 intracellular activity in novel and familiar environments. Elife 6, e23040 (2017).

[51] Malvache, A., Reichinnek, S., Villette, V., Haimerl, C. & Cossart, R. Awake hippocampal reactivations project onto orthogonal neuronal assemblies. Science 353, 1280–1283 (2016).

[52] Simpson, A. J. & Fitter, M. J. What is the best index of detectability? Psychological Bulletin 80, 481 (1973).

